# Quantitative assessment of pain threshold induced by a single-pulse transcranial magnetic stimulation

**DOI:** 10.1101/2020.02.13.946921

**Authors:** Keisuke Tani, Akimasa Hirata, Satoshi Tanaka

**Author notes:** Corresponding author Satoshi Tanaka.

## Abstract

**Objective:** Transcranial magnetic stimulation (TMS) is commonly used in basic research to evaluate human brain function. Although scalp pain is a side effect, no studies have quantitatively assessed the TMS intensity threshold for inducing pain and whether sensitivity to TMS-induced pain differs between sexes.

**Methods:** We measured pain thresholds when single-pulse TMS was applied over either Broca’s area (BA) or left primary motor cortex (M1). We compared these thresholds with motor threshold for inducing motor evoked potential (MEP) through M1 stimulation. We also compared pain thresholds for BA and M1 between males and females.

**Results:** Pain thresholds for both sites were significantly lower than motor threshold. Further, the pain threshold for BA was much lower than that for M1. No significant difference was observed between sexes.

**Conclusion:** The results suggest that TMS at an intensity equivalent to motor thresholds, which is often used in experimental or clinical studies, causes slight scalp pain.

**Significance:** Experimental designs using TMS to evaluate functional relationships between brain and behaviors should consider scalp pain and reduce its likelihood as much as possible.

**Highlights:**

- We investigated pain thresholds induced by a single-pulse TMS over the head.
- Pain thresholds for TMS over Broca’s area (BA) and primary motor cortex (M1) were much lower than motor threshold.
- No significant differences in the pain thresholds were observed between sexes.

## 1. Introduction

Transcranial magnetic stimulation (TMS) is widely used in basic research as a tool for evaluating human brain function. Additionally, several studies have demonstrated the effectiveness of repetitive TMS on the recovery of motor, cognitive, or mental function in patients with neurological or psychiatric disorders (e.g. Khedr et al., 2010; Ren et al., 2014; Kobayashi and Pascual-Leone, 2003). In TMS, an electric current in the brain is induced by producing a strong magnetic field via current flowing the coils, which can lead a temporary change in brain activity (Rossi et al., 2009). TMS also causes head and face muscles to contract and stimulates cutaneous fibers, which often leads to pain or discomfort on the scalp (Rossi et al., 2009; Wassermann, 1998; Rumi et al., 2005). This side-effect of TMS influences aspects of task performance such as accuracy and reaction time (Abler et al., 2005; Meteyard and Holmes, 2018), and to interfere with successful completion of experiments (Wassermann, 1998; Satow et al., 2002). Even though researchers have assessed the degree of pain and the area in which pain is perceived during TMS of certain intensities (Arana et al., 2008; Meteyard and Holmes, 2018), to our knowledge, no studies have quantitatively evaluated what TMS intensities actually cause pain (i.e., the pain threshold).

In most neurophysiological and neuropsychological studies, TMS intensity is determined based on the motor threshold, which is defined as the minimum current intensity of TMS that produces a pre-defined motor-evoked potential (MEP) amplitude in a target muscle (Rossini et al., 1994). Indeed, an intensity equivalent to 100%–120% of motor threshold is often applied in these types of experiments (e.g., Wu et al., 2000; Lazzaro et al., 2002). Thus, evaluating whether the pain threshold is higher or lower than motor threshold is critical. The primary purpose of the present study was to measure pain thresholds when single-pulse TMS is delivered to a motor center (primary motor cortex; M1) or a speech motor center (Broca’s area; BA), and compared them with motor threshold.

While several studies have demonstrated that females have a higher sensitivity to experimental pain stimuli than males (Riley et al., 1998; Andersen et al., 2015), others have showed that pain thresholds do not differ between sexes (Isselee et al., 1998; Racine et al., 2012). In their meta-analysis, Riley et al. (1998) suggested that a large number of participants are necessary to test a sex difference in pain thresholds. In this study, we therefore assessed whether pain thresholds for TMS differed between males and females using relatively large sample size.

## 2. Methods

### 2.1. Participants

The study was approved by the ethical committee of Hamamatsu University School of Medicine and was in accordance with the Declaration of Helsinki. A meta-Analysis (Riley et al., 1998) has shown that 41 participants per group are necessary to assess the sex differences in pain threshold with an enough power (d = 0.70). Based on this, 82 healthy individuals (41 males and 41 females; age: 26.4 ± 12.3 years) were recruited for the study. All provided written informed consent before the experiment. We confirmed through questionnaires that no participants had epilepsy, none had a family history of epilepsy, and none had any neurological or psychiatric disorders. The protocol of this study was pre-registered in the University Hospital Medical Information Network (UMIN) clinical registry (registration number: 000029783) in Japan.

### 2.2. TMS device

We delivered single-pulse TMS to the scalp of each participant using a Magstim stimulator (Magstim, 200, Magstim Co. Ltd, UK) with a figure-eight coil (70 mm diameter; Magstim Co. Ltd, UK).

### 2.3. Stimulation site and coil orientation

We applied TMS over the targeted brain locations using neuro-navigation. Before the TMS experiment, each participant underwent a T1-weighted magnetic resonance imaging (MRI) head scan with a 3T scanner (Discovery MR750 3.0T, GE Healthcare Japan, Japan). The scanner setting was as follows: repetition time (TR) = 7.2 ms, echo time (TE) = 2.1 ms, flip angle (FA) = 15°, field of view (FOV) = 25.6 cm^2^, voxel size = 1 mm × 1 mm × 1 mm, and matrix = 256 × 256. Based on the MRI images, we created a 3D cortical surface model of each participant using a frameless navigation system (Brainsight, Rogue Research Inc, Canada). Using this device, we could continuously monitor the position and orientation of a TMS coil relative to participant’s head by capturing the reflection markers mounted on the head and the coil with a camera. This allowed us to accurately stimulate the regions of interest (left M1 or BA) during the experiment.

The stimulation locations were anatomically identified. For M1, we used the center of the “hand knob”, which is a landmark of hand motor cortex (Yousry et al., 1997). This method has been shown to be as reliable as the standard method of functionally determining the stimulation site based on the amplitude of induced MEPs (Sparing et al., 2008). For BA, we used Brodmann area 44. The orientation of the magnetic coil for M1 was set as a posterior and lateral handle orientation, 45 degrees relative to the antero-posterior axis of the head, which appears to be the optimal angle for inducing MEPs (Mills et al., 1992). The coil orientation for BA was set as a posterior handle orientation, i.e. parallel to antero-posterior axis of the head, seen from the lateral side.

### 2.4. Threshold measurements

For each participant, we assessed the pain thresholds for M1 and BA and the motor threshold for M1. During measurements, participants sat on a reclining chair and were asked to relax. The order of measurements (stimulation site or threshold) was randomized across participants.

#### 2.4.1 Pain threshold

Participants were asked to verbally report the presence or absence of scalp pain every after each stimulation. We measured pain thresholds using an adaptive staircase method; we decreased the intensity when the participant reported pain, and increased the intensity when they reported the absence of pain. Pain thresholds were defined as the minimum intensity that induced pain in at least 5 of 10 trials; the minimum intensity is represented in terms of percentage of maximum stimulator output (MSO).

#### 2.4.2. Motor threshold

We measured the peak-to-peak amplitude of MEPs induced by TMS over left M1 (the hand knob). Electromyography (EMG) of the right first dorsal interosseous muscle (FDI) was recorded using Ag/AgCL surface electrodes (10 mm in diameter). The EMG signals were amplified, bandpass filtered between 16–470 Hz, and sampled at 3 kHz by a Rogue EMG device. During this measurement, participants were asked to keep their hands as relaxed as possible. A researcher carefully confirmed whether the muscles were relaxed by watching the EMG activity on a monitor when applying each stimulation. Similar to the pain threshold, we used an adaptive staircase method and defined the motor threshold as the minimum intensity that induced MEPs whose amplitudes were larger than 50 μV in at least 5 of 10 trials.

### 2.5. Data analysis

Pain and motor thresholds were determined for each participant. Shapiro-Wilk tests revealed that threshold values were not normally distributed across participants (*p* < 0.001 for BA, *p* = 0.005 for M1). Therefore, we used a Friedman test to compare the three thresholds, followed by post-hoc Scheffé tests. Additionally, we compared pain sensation for BA and M1 between males and females using Mann-Whitney U tests.

## 3. Results

The results for pain and motor thresholds are presented in Figure 1A. Median (1^st^, 3^rd^ quartiles) pain thresholds for BA and M1 were 24.0% (32.8, 19.3) and 43.0% (51.0, 36.0) of MSOs, and motor thresholds were 54.5% (64.0, 46.3). Minimum value of pain thresholds for BA and M1 were 10% and 24%, respectively. A Friedman test revealed a significant difference between the thresholds (χ^2^_(2)_ = 112.2, *p* < 0.001). Post-hoc tests showed that pain thresholds for BA were significantly lower than motor and pain thresholds for M1 (both *p* < 0.001), and that the pain thresholds for M1 were significantly lower than the motor thresholds (*p* < 0.001). Pain thresholds for BA and M1 were lower than motor threshold in 78 (95%) and 67 participants (82%), respectively.

**Figure 1.**
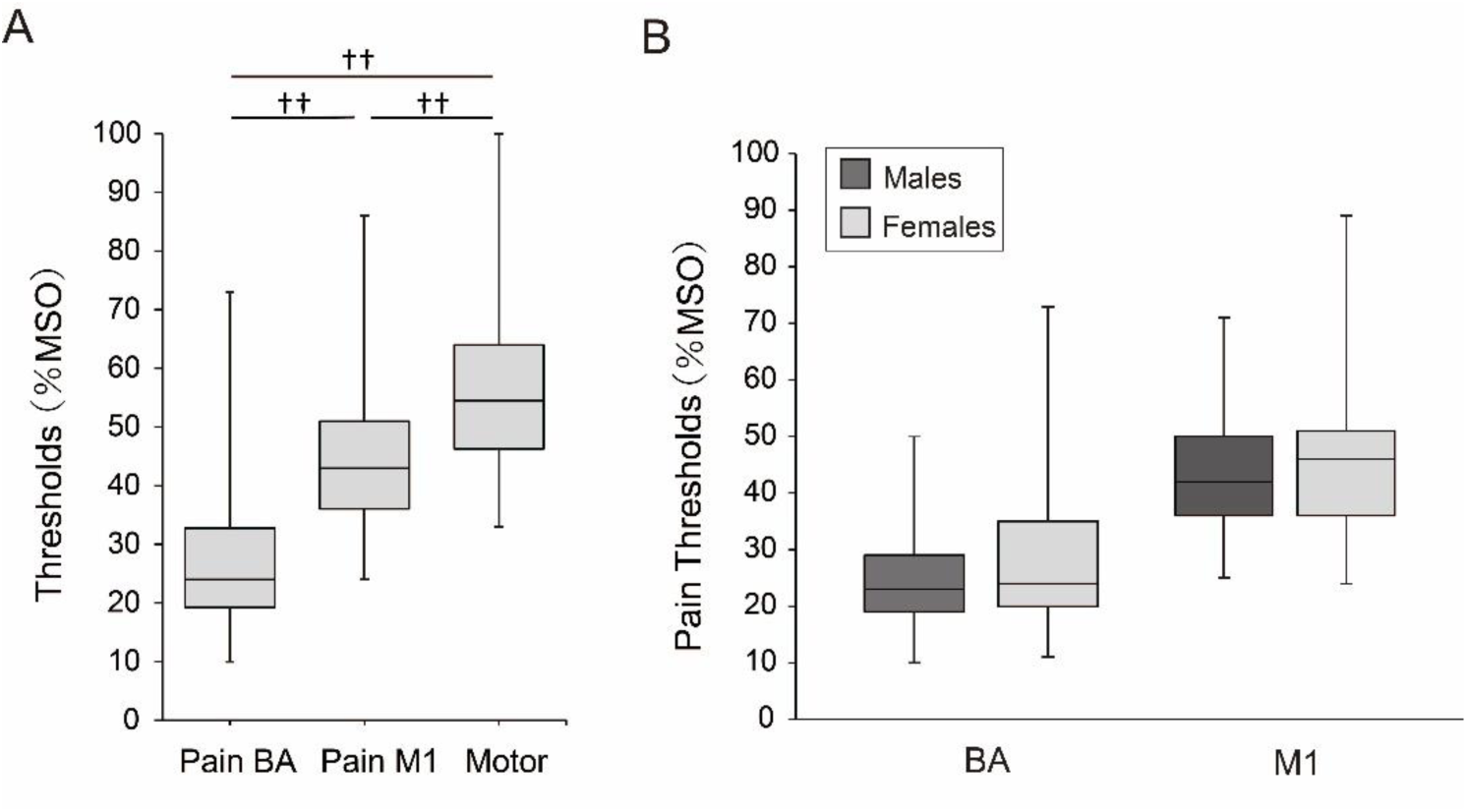
**A.** Box-whisker plots for pain and motor thresholds. Each box covers the range between the 1st and 3rd quartiles of each threshold. The horizontal line within the box represents the median value for each threshold. Upper and lower ends of each whisker represent the maximum and minimum values for each threshold, respectively. ^††^, *p* < 0.001 **B.** Box-whisker plots for pain thresholds for BA and M1 for males and females.

Figure 1B compares pain thresholds for males and females. Median (1^st^, 3^rd^ quartiles) pain thresholds for males and females were 23.0% (29.0, 19.0) and 24.0% (35.0, 20.0) for BA, and 42.0% (50.0, 36.0) and 46.0% (51.0, 36.0) for M1, respectively. Mann-Whitney U tests revealed no significant differences between sexes both for either BA (z = 0.79, *p* = 0.43) or M1 (z = 0.60, *p* = 0.55).

## 4. Discussion

In most clinical or experimental studies using TMS, the intensity of stimulation is determined as equivalent to or above individual motor thresholds, but whether stimulation at this intensity causes scalp pain is unclear. The present study quantitatively evaluated pain thresholds for single-plus TMS delivered to M1 and BA, and compared them with motor threshold. We found that on average, the pain thresholds for both BA and M1 were much lower than the motor threshold, and that the pain thresholds for both sites were lower than the motor threshold in more than 80% of participants. These results indicate that participants feel slight pain when TMS is applied at intensities equivalent to the motor thresholds. It has been demonstrated that pain or discomfort induced by TMS influences task performance (Abler et al. 2005; Meteyard and Holmes, 2018; Holmes and Meteyard, 2018). For instance, Abler et al. (2005) showed that the error rate on a visual memory task increased with the magnitude of subjective discomfort induced by TMS. Based in these findings, we must consider that any temporary changes in motor or cognitive performance observed after applying TMS to a particular brain region might also be related to the side effect of pain. Therefore, to properly evaluate the functional relationships between brain and behavior using TMS, we need to design experiments in which the influence of pain sensation is controlled for. This could take the form of a sham condition in which the same degrees of pain sensation is caused on the scalp.

We also found that pain thresholds were much lower for BA than for M1. This difference in pain threshold between sites could be related to anatomical differences in muscle volume. While the tissues of temporalis muscle are thickly distributed on the scalp above BA, aside from the epicranial aponeurosis, muscle tissues are not distributed above M1 (Netter, 1989). Based on this fact, the present result proposes that TMS-induced pain can be attributed primarily to muscle twitches rather than activation of cutaneous nerves. Indeed, some studies have suggested that magnetic stimulation can stimulate the peripheral or central neuromuscular system without strongly activating skin nociceptors (Barker et al., 1987; Barker, 1991; Han et al., 2006). TMS over BA would strongly stimulate the muscle nociceptors (i.e., free nerve endings of Aδ or C fibers), which would result in easy elicitation of pain.

We also evaluated differences in pain thresholds between males and females with an appropriate sample size (Riely et al., 1998). In contrast to previous findings that pain thresholds in females were lower than that in males (e.g. Riley et al., 1998; Andersen et al., 2015), we observed no significant sex differences. One reason could be that sex differences in pain thresholds depend on the type of pain. Indeed, higher sensitivity in females has been consistently demonstrated for pressure pain (Andersen et al., 2015) and electrical pain (Riley et al., 1998), but not for other modalities such as ischemic, cold, or heat pain (Racine et al., 2012). It is likely that TMS-induced pain is also derived from a mechanism different from pressure or electrical pain, which would result in no significant sex differences in pain thresholds for TMS.

In the present study, we only subjectively measured pain thresholds. The details of how TMS stimulates each tissue within the scalp (skin, fat, or muscle) remains unclear. Thus, the present result cannot exclude the possibility that the activation of peripheral nerves also contributes to pain sensation. Skin thickness seems to vary depending on cranial sites (Young, 1959), which might partially explain the differences in pain thresholds between the cortical sites (Rashed et al. 2019). To clarify the mechanism of pain sensation, further computational studies are required to estimate the distribution of the electric field over skin and muscle tissues induced by TMS.

## 5. Conclusion

The present study shows TMS induces pain at intensities equivalent to motor threshold especially over the BA, although its sensation is relatively weak around pain thresholds. Therefore, TMS researchers need to remind themselves that some participants could feel pain even when TMS intensity is lower than the motor thresholds in their experiments.

## Acknowledgements

This study was supported by Ministry of Internal Affairs and Communications, Japan. We thank Adam Phillips, PhD, from Edanz Group (https://en-author-services.edanzgroup.com/) for editing a draft of this manuscript.

